# Suppression of miRs-497/195 axis possibly confers endocrine therapy resistance via elevated expression of FLT4 and the noncoding RNA MIR503HG

**DOI:** 10.1101/2024.05.14.594132

**Authors:** Saheli Pramanik, Partha Das, Monalisa Mukherjee, Kartiki V. Desai

**Affiliations:** Biotechnology Research Innovation Council-National Institute of Biomedical Genomics (BRIC-NIBMG), Kalyani, India; Ph.D. Program, Regional Centre for Biotechnology, 3rd Milestone, Faridabad-Gurugram Expressway, Faridabad, India

**Keywords:** ceRNA, Tamoxifen, LTED, JMJD6, CNC network, Endocrine therapy resistance

## Abstract

**Background:** Endocrine therapy resistance (ETR) in breast cancer is achieved via multiple pathways including a decrease in ER, dysregulation of cell cycle genes, and/or mutations in ER/co-activators/co-repressors. We have reported earlier that high expression of Jumonji domaining containing protein 6 (JMJD6) induced ETR by depleting ER expression. In this study, 3 cellular models representing distinct ETR pathways; Tamoxifen resistant (TAMR), Long-term Estrogen deprived (LTEDI), JMJD6 overexpressing (JOE) cells, and parental MCF7 were subjected to RNA-sequencing, CNC, and ceRNA network analysis. We hypothesised that post-comparison RNA regulations that are common to all cell lines, will reveal actionable markers and targets. These will be shared by all patients with ET-resistant disease, independent of the initiating event.

**Results:** 170 differentially expressed genes were found, of these, 73 maintained the same directionality in expression (ETR cassette genes). These genes segregated TCGA ER+ tumors into two groups, one intermixing with ER-tumors. Pathway-based curation of ETR genes identified 21 genes (7 up- and 14 down-regulated) that participated in multiple cancer hallmark pathways. Genes upregulated in ETR cells were less expressed in ER+ tumors at diagnosis when compared to normal breast samples but their higher expression indicated adverse survival outcomes. Next, these genes were used for CNC and ceRNA network construction and a triad FLT4:MIR503HG:miR-497/195/424 was discovered. The expression levels of miRNAs were predicted via network analysis and quantitative RT-PCR was used to validate the down regulation of miR-497/195/424 and upregulation of their targets, FLT4 and MIR503HG in ETR cells.

**Conclusions:** We show that total RNA-seq data can be successfully used to predict actionable miRNAs that achieve drug resistance. Re-expression of ETR genes such as FLT4 in tumor cells, that are less expressed at diagnosis, may be indicative of ETR onset. Finally, ETR may arise due to suppression of miR-424/497/195 leading to higher expression of FLT4 and MIR503HG. We posit that FLT4 may be a suitable target and RT-PCR analysis of this RNA triad could be developed as a detection strategy for ETR in ER+ breast cancer.

## 1. INTRODUCTION

Breast cancer (BrCa) is the most common cancer in women and has high mortality rates due to high cellular and molecular heterogeneity (1,2). 70% of diagnosed breast cancer is estrogen receptor positive (ER+) and is treated with endocrine therapy (ET) in the adjuvant setting. Selective Estrogen Receptor Modulators (SERMs) such as Tamoxifen (TAM), Selective Estrogen Receptor Degraders (SERDs) like Fulvestrant and Aromatase Inhibitors (AIs) Letrozole, and Anastrozole have successfully been used as the first line of therapy (3). However, about 30-40% of women recur and become non-responsive to endocrine therapy (4). Current treatment options for these women are very limited and ET resistance (ETR) remains a major challenge for the treatment of both recurrent and metastatic breast cancer (MBC). Development of tests that forecast ETR and identify novel targets for treating resistant disease is a major area of research.

Several genetic tests such as Oncotype Dx, EndoPredict, and MammaPrint provide the risk of recurrence scores and many studies have developed gene signatures associated with TAM, Fulvestrant, and ETR in breast cells and cell line xenografts via transcriptome analysis (5–9). Few of these signatures have been adapted in the clinic to estimate ETR because rarely do tumors display inherent resistance. Such resistance is acquired during the course of treatment and transcriptomic profiles during treatment from repeated biopsies are unavailable. Using cell line and patient sample data, pathways leading to such resistance are well worked out and extensively described in the literature (10). Molecular heterogeneity, differential response and outcomes to the same therapy, and use of complex pathways in ETR, limit the use of discovered gene signatures in the clinic. Broadly, events leading to ETR can be grouped into 3 types-1) lowering of ER expression, 2) acquired resistance by ligand-independent activation of ER and/or high expression of cell cycle genes, or epigenetic changes, and 3) by gain of activating mutations in ER itself. Different modes of ETR involve distinct factors that elicit different gene expression programs.

We propose that an integrated RNA network analysis to find common genes/pathways using cell lines representing various forms of ETR could be useful in designing biomarkers that indicate potential ETR, independent of the initiating event. It is clear that the expression of not only the coding mRNAs but also the non-coding genome influences ETR (11). While the long non-coding RNAs (lncRNAs) regulate mRNA expression by cis-/trans-regulation at the transcriptional level, microRNAs (miRNAs) work post-transcriptionally to interfere with the translation of the protein encoded by its target mRNA (12). Often the same miRNA has sequences complementary to both lncRNAs and mRNAs. Various regulatory outcomes are possible based on the levels of miRNA, lncRNA, and mRNAs present at any given time (13). Clearly, the ratio of mRNA:lncRNA:miRNA will tend to determine the final expression levels of a gene/protein. Typically, co-expression and competing endogenous RNA networks are used to understand the complex RNA regulation and they have been successfully used to segregate ER+ and ER-tumors, understand the response to Tamoxifen in cancer patients, and generate lncRNA or miRNA signatures (11,14–17). ceRNA networks for Tam sensitive and TAMR cells have been constructed previously and MCF7 derived LLC3 and LCC9 Tam resistant cells (18,19). ceRNA networks for alternate modes of ETR have not been constructed earlier.

Here we generate the total transcriptomes of ETR cell line models. MCF7 derived Tamoxifen resistant (TAMR), Long-Term Estrogen Deprived-Independent (LTEDI) that do not require estrogen for proliferation (20), and JOE cells that overexpress Jumonji Domain Containing protein 6 (JMJD6). Our laboratory has developed TAMR cells that have high ER, RET, and phospho-AKT levels, LTEDI cells having high ER, ERK1/ERK2 expression whereas JOE cells harboring low ER, high RET, and high ERK1/ERK2 levels. JOE cells are insensitive to endocrine therapy and re-sensitized is possible by depleting JMJD6 (21). Incidentally, TAMR and LTED cells also showed high expression of JMJD6, and depletion of JMJD6 in TAMR cells restored sensitivity to TAM (21). JMJD6 is a major epigenetic regulator of gene expression, and induced cyclin E2, E2F targets, and G2/M checkpoint genes to promote cell cycle advancement in breast cancer cells, thus removing their dependency on ER (21–23). Increased cyclin E2 was also identified as a marker of prognosis and persistent Fulvestrant resistance in MBC (24). Clearly, each cell model system can evade TAM induced growth restriction via overlapping and distinct modes of ETR. In this study, genes common and similarly regulated in all 3 cell lines were identified, filtered by pathway-based curation, and subjected to CNC and ceRNA network analysis.

## 2. MATERIALS AND METHODS

### 2.1. Cell culture

MCF7 cells were purchased from the American Type Culture Collection (ATCC, VA, USA). Cells were cultured in Dulbecco’s modified Eagle’s medium (DMEM) (GIBCO, USA) with 5% fetal bovine serum (FBS) (GIBCO, USA) and 1% PenStrep (GIBCO, USA) in humidified 5% CO_2_ incubator at 37°C. Cell lines were negative for mycoplasma presence (Lonza, Switzerland). Generation of JMJD6 overexpression (JOE) cells, TAMR, and LTEDI cells are described previously (21).

### 2.2. Cell Viability and Colony Forming Assay

Cell viability assay was performed by CCK8 kit (Sigma Aldrich, USA) according to the manufacturer’s protocol. MCF7 cells were starved for 3 days in estrogen-deprived media. After 3 days, MCF7 and LTEDI cells were plated at a density of 10^4^ cells/well in 96 well plate. Treatments were 1μM 4-hydroxy TAM (4-OHT) and ethanol (as vehicle control) for 72 hours. Absorbance was measured at 450 nm. LTEDI cells (10^3^ cells/well) were seeded in 6-well plates. The next day, cells were treated with ethanol/1 μM 4-OHT for 9 days with a daily media change. Cells were fixed with 4% paraformaldehyde and stained using a crystal violet solution. The number of colonies was quantified and cells were photographed.

### 2.3. RNA-sequencing, Library Preparation and Data Analysis

RNA was isolated from MCF7, TAMR, and LTEDI cells by using a Qiagen RNAeasy mini kit (Qiagen, Germany). Library preparation was carried out at the NIBMG Core facility for 100 bp paired-end reads (75 million each) using NOVASeq. Following FASTQC quality check FAST Q files were trimmed with CutAdapt to remove the adapter sequences and aligned with the human genome (GrCh38.1) by HiSAT2.0 algorithm. Alignments were processed by the FeatureCounts package for obtaining RNA/ transcript counts for differentially expressed coding (DEGs) and long non-coding RNAs (DELs) separately. Data is available at GEO Omnibus (GSE255704). Limma-voom analysis followed by Benjamini-Hochberg correction was used to obtain DEGs/DELs significant at adjusted p-value ≤ 0.05 and log fold change= 1(log2FC) (https://usegalaxy.org). Cluster 3.0 and Tree View programs were used for heatmap generation and visualization. Pathway enrichment analysis was done by EnrichR (https://maayanlab.cloud/Enrichr/). Top 10 significant pathways are shown for further comparison. Chord diagrams for genes appearing in various pathways was constructed using SRplot. To obtain DEG data from Fulvestrant resistant MCF7 cells, GSE118713 was analysed using GEO2R and our internal pipeline (25).

### 2.4. Co-expression (CNC) and ceRNA networks

To associate the lncRNAs with target mRNAs, we constructed the lncRNA-mRNA co-expression network. For each pair of DEGs and DELs, the Pearson correlation was calculated and those pairs with correlations of 0.98 or greater were used to construct the network. For cis-regulation, the genomic distance between each pair was calculated using data obtained from Biomart (https://www.ensembl.org/biomart/martview/). None of the pairs were located within 10 kb (data not shown). For the ceRNA networks (mRNA-lncRNA-miRNA), first, the miRNAs for DELs were predicted from the miRcode (http://www.mircode.org) and both miRcode and TargetScan (v7.2) (https://www.targetscan.org) database were used for miRNAs and DEGs. A list of mRNA-lncRNA-miRNAs common to both TAMR-LTED was obtained. JOE networks were constructed separately. Pathway-based curation was carried out to obtain DEGs that appeared in multiple significant pathways to give 7 upregulated and 14 downregulated genes (21 genes). The mRNA-lncRNA-miRNA interaction networks were constructed using 21 DEGs and corresponding lncRNAs and miRNAs. Cytoscape (https://cytoscape.org, version 3.10.0) was used to construct and visualize all networks.

### 2.5. RNA extraction and quantitative real-time PCR (qRT-PCR)

Quantitative real-time PCR (qRT-PCR), using primers described in Supplementary Table 1 was carried out as described earlier (21). C_T_ values of gene-specific primers were normalized to the C_T_ value of β-actin. Fold change in gene expression across samples was calculated by the formula 2 ^(-ΔΔCT)^ using values from ‘Vec’ or MCF7 cells converted to fold change 1.

### 2.6. Stem-loop PCR for miRNA

Three miRNAs (miR-497, miRNA-424, miRNA-195) were selected for qRT-PCR. Total RNA was extracted from all the cell lines using RNA extraction kit (AllPrep DNA/RNA/miRNA Universal Kit, QIAGEN). One microgram of total RNA was reverse-transcribed using each miRNA-specific stem-loop primer (Supplementary table 1). Expression of each miRNA was quantified by using SYBR green mix (KAPA BIOSYSTEMS, South Africa). C_T_ values of miRNAs were normalized with C_T_ value of U6. Fold change in gene expression across samples was calculated by the formula 2 ^(-ΔΔC)^ using values from ‘Vec’ or MCF7 cells converted to fold change 1.

### 2.7. TCGA Data and Survival Analysis

cBioPortal and GDC portal were used to download the Pan-Cancer Atlas breast cancer related dataset counts with clinical data, age, subtype information, inclusion-exclusion criteria, etc of total sample 1084 ( LumA 499, LumB 197, Basal 171, Her2 78, Normal like 36, no data-103) and had follow-up data (https://www.cbioportal.org/study/clinicalData?id=brca_tcga_pan_can_atlas_2018). mRNA expression z-scores relative to all selected samples were used to determine gene expression patterns of select genes. The relative expression of selected genes as tumor versus normal was plotted by UALCAN (https://ualcan.path.uab.edu). The median expression value was used as a cut-off and significant difference between groups was assessed using the unpaired two-tailed Student’s t-test. The prognostic impact of selected genes was analysed using the Kaplan-Meier Plotter (https://www.kmplot.com/analysis/).

## 3. RESULTS

### 3.1. Effect of Tamoxifen on LTEDI cells

Both agonistic and antagonistic activity of TAM has been reported for LTEDI cells in the literature, however, independence from the requirement of exogenous E_2_ for proliferation is common to all LTEDI cells (19). To determine the nature of our cells, we studied the effect of 4-OHT both in short-term cultures using CCK-8 assays and in longer-term cultures by colony forming assay. While parental MCF7 cells desist, LTEDI cells continue to proliferate in the presence of 4-OHT over 72 hours (Figure 1A). Secondly, the overall proliferation rate of LTEDI is higher than parental MCF7 cells in the absence of 4-OHT, and this is further enhanced by the addition of the drug (Figure 1B). Our data indicates that 4-OHT may serve as an agonist in LTEDI cells.

**Figure 1.**
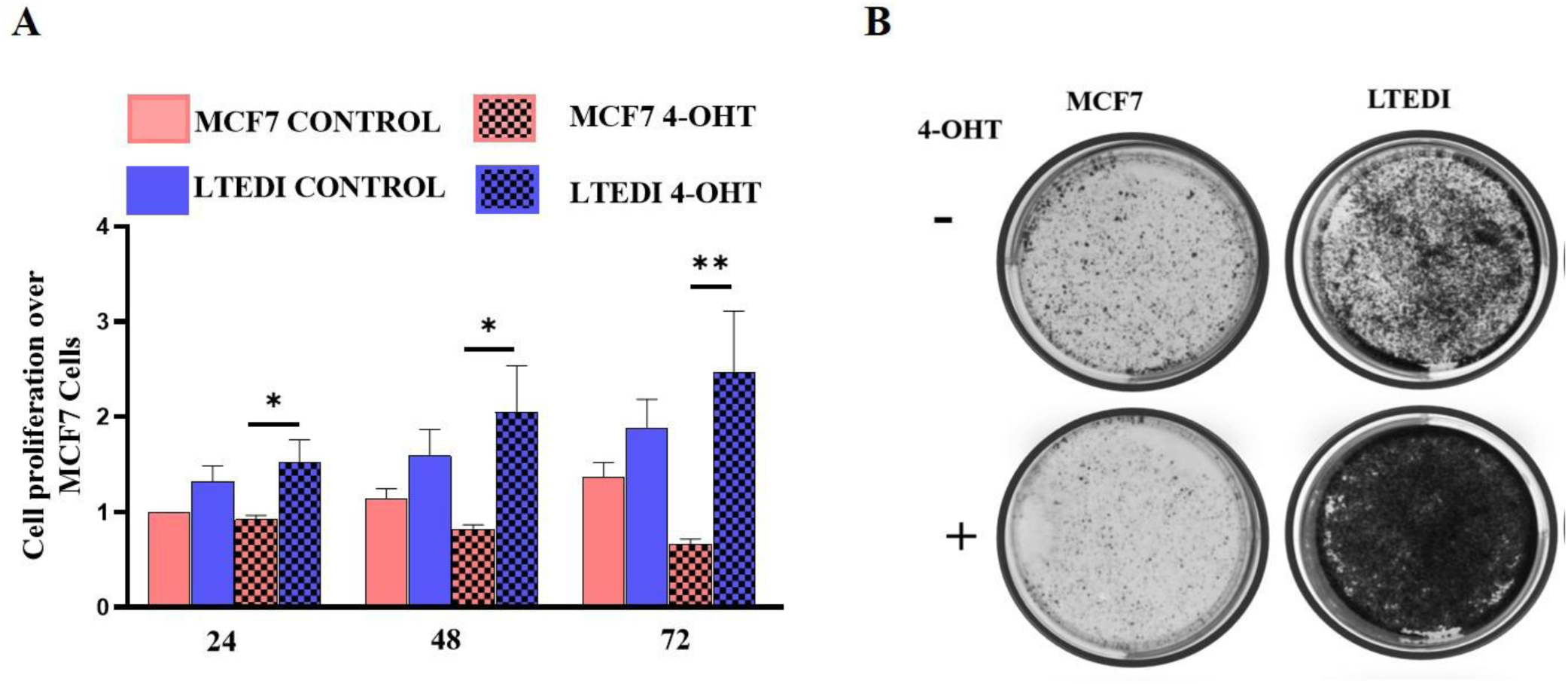
Effect of 4-OHT in LTEDI cells. **(A)** Cell proliferation assay of MCF7 and LTEDI cells in the presence and absence of 4-OHT measured by CCK8 kit. \**p* ≤ 0.05, \*\**p* ≤ 0.01, and \*\*\**p* ≤ 0.001. **(B)** Colony Forming Assay of TAMR and LTEDI cells in the presence and absence of 4-OHT.

### 3.2. Identification of commonly regulated DEGs in JOE, TAMR and LTEDI cells

The workflow for identifying mRNA, lncRNA, and miRNAs of our analysis is shown in figure 2. We published the transcriptomic program of JOE cells expressing low ER earlier and identified about 1100 genes that were differentially expressed in these cells (21). RNA-seq analysis of TAMR cells identified 8685 genes as significantly differentially expressed (adjusted p-value ≤ 0.05) when compared to parental MCF7 cells. 1609 genes were upregulated and 2332 genes were down-regulated (log_2_FC=1) (Figure 3 A, B supplementary file 1). In LTEDI cells, 5630 DEGs were found (adjusted p-value ≤ 0.05), of which, 687 genes were upregulated and 1476 genes were down-regulated (log2FC=1) (Figure 3 A, B supplementary file 1). Pathway enrichment analysis (EnrichR) was used to compare the two cell line models and significant pathways are shown (Adjusted p-value ≤ 0.05, Figure 3 C). Estrogen response early, Estrogen response late, and Interferon alpha and gamma pathways were regulated in both TAMR and LTEDI cells, however, the number of genes was variable. These are indicated by the size and color variation in the bubble plot (Figure 3 C). mTORC1 signalling, p53 pathway, Hypoxia, Glycolysis, Cholesterol homeostasis, Xenobiotic metabolism, Epithelial-Mesenchymal Transition (EMT) were unique to TAMR cells while TNF-alpha signalling via NF-kB pathways, KRAS signalling Up, and Wnt-beta Catenin signalling was found in LTEDI cells (Figure 3 C).

**Figure 2:**
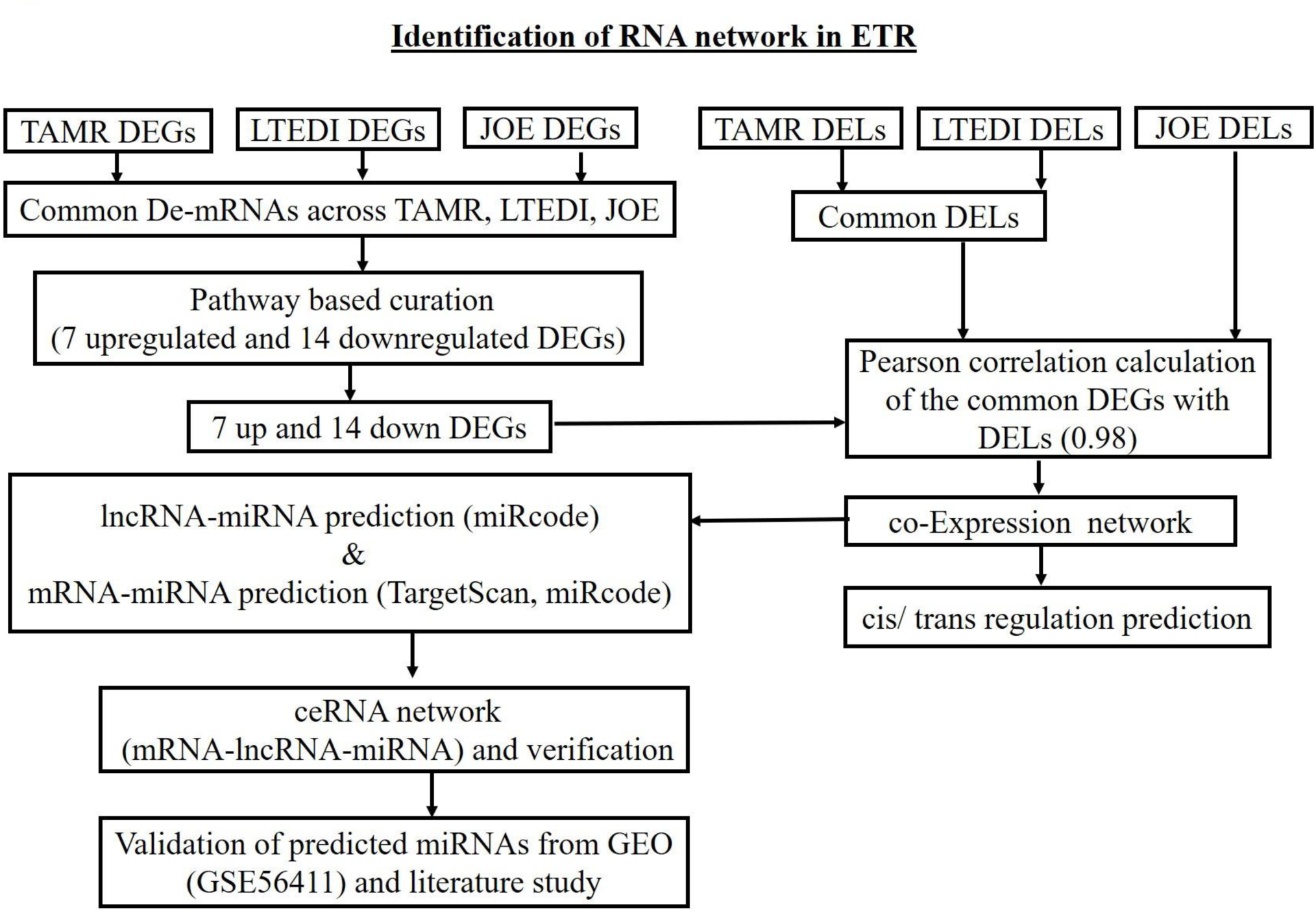
Workflow for constructing the RNA regulatory networks.

**Figure 3:**
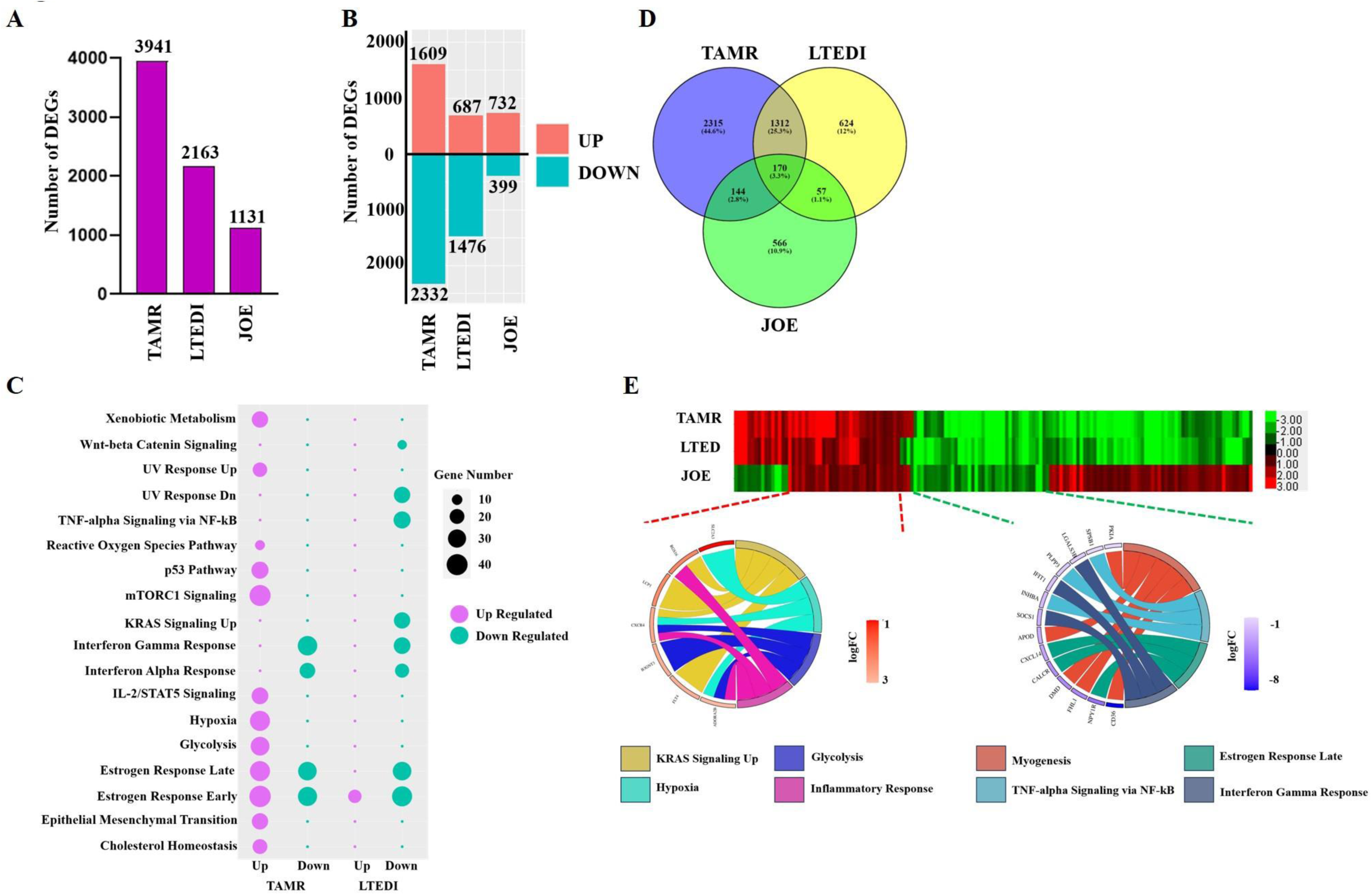
Transcriptome analysis of cell lines. **(A)** DEGs were found in TAMR, LTEDI, and JOE cell lines compared to MCF7 cells. Numbers indicate DEGs in respective cell lines. **(B)** Upregulated and downregulated DEGs in TAMR and LTEDI cell lines. **(C)** Bubble plot showing significant pathways found in TAMR and LTEDI cell lines. The X-axis shows the number of genes in each pathway and Y-axis indicates the name of the pathway. The pathways and bubble size key are indicated on the right-hand side. **(D)** Venn diagram of the intersection of TAMR, LTEDI, and JOE DEGs. **(E)** Heatmap of 170 genes with green (downregulated) to red (upregulated) DEGs. A Chord diagram showing pathway curated genes (7 upregulated and 14 downregulated) in multiple pathways. Significant pathways are shown on the right, and the fold change of core genes is shown on the left. Left–right connections indicate gene membership in a pathway’s leading-edge subset.

Our goal was to obtain genes that accounted for an escape from drug induced growth restriction. The intersection of TAMR and LTEDI DEGs yielded 1482 genes. Interestingly 94% of these genes showed the same directionality in regulation; with 453 induced and 973 suppressed DEGs. TAMR-LTEDI-JOE intersection had 170 genes common amongst them, with 73 genes sharing the directionality (all up/all down/all cells) and referred to as the endocrine therapy resistance cassette genes (ETR cassette) (Figure 3 D). Out of 73 genes, 33 were upregulated and 40 downregulated (Figure 3 E). The remaining 97 genes out of 170 genes were regulated in the opposite direction in JOE cells and mostly contained the ER regulated genes. In JOE, ER levels are suppressed but ER is robustly expressed in both TAMR and LTEDI cells, which may partially account for this observation. Since Fulvestrant is used for endocrine therapy, we carried out a similar analysis (both by GEO2R and our pipeline) in Fulvestrant resistant MCF7 cell data described in GSE118713 (25). After comparison with TAMR-LTEDI-JOE data, we found 85 genes were common (Supplementary figure 1 A and B), 26 upregulated and 11 downregulated genes.

The ETR cassette was enriched for KRAS signalling, Hypoxia, Glycolysis, Inflammatory response pathways in the upregulated genes and Myogenesis, TNF-alpha signalling via NFkB, Estrogen response late, Interferon gamma response represented in the downregulated genes. We noticed that 7 induced DEGs and 14 downregulated DEGs participated in every significant pathway identified, indicating their importance in ETR (Figure 3 E). Next, we assayed the expression of the ETR cassette in TCGA data and used pathway-based curated genes (7up:14down) to construct CNC and ceRNA networks.

### 3.3. Gene expression in breast cancer patient samples

In the publicly available pan cancer (TCGA breast) dataset, we analysed the Dead/Alive clinical status and ER+ subtype specific gene expression data using death as an outcome to represent a poor response to treatment. We failed to obtain DEGs, but this was expected since ET resistance is acquired over prolonged exposure to drugs and long-term estrogen deprivation, and it is rarely intrinsic. However, a supervised hierarchical clustering analysis of 468 samples (113 normal, 356 Luminal A and B) using mRNA expression z-scores relative to all samples showed that ETR cassette genes effectively segregated tumors from normal samples (data not shown). Since very little data comparing longitudinally collected samples at diagnosis and following recurrence is available, we tested if our ETR cassette was better at discriminating the ER+ tumors at diagnosis. As this set originated from cell models that had an E_2_-ER independent axis of growth, we hypothesized that some ER+ tumors with the potential to avoid ET would cluster better with ER-tumors. Supervised hierarchical clustering analysis of 1084 samples (all subtypes) using mRNA expression z-scores relative to all samples (log RNA Seq. V2 RSEM) for 73-gene ETR cassette was carried out and is shown in Figure 4 A (supplementary figure 2 A). Interestingly, the 33 upregulated ETR split the ER+ subtype into two groups, one in which ER+ tumors were mixed with TNBCs, and another that remained distinct from ER-tumors. However, the down-regulated genes showed no consistent pattern of expression (Supplementary figure 2 A). This indicates that this 33 gene ETR set obtained from cellular models of ET resistance may be useful in the segregation of ER+ tumors into those that respond versus those that do not. However, more studies are required to validate this observation.

**Figure 4:**
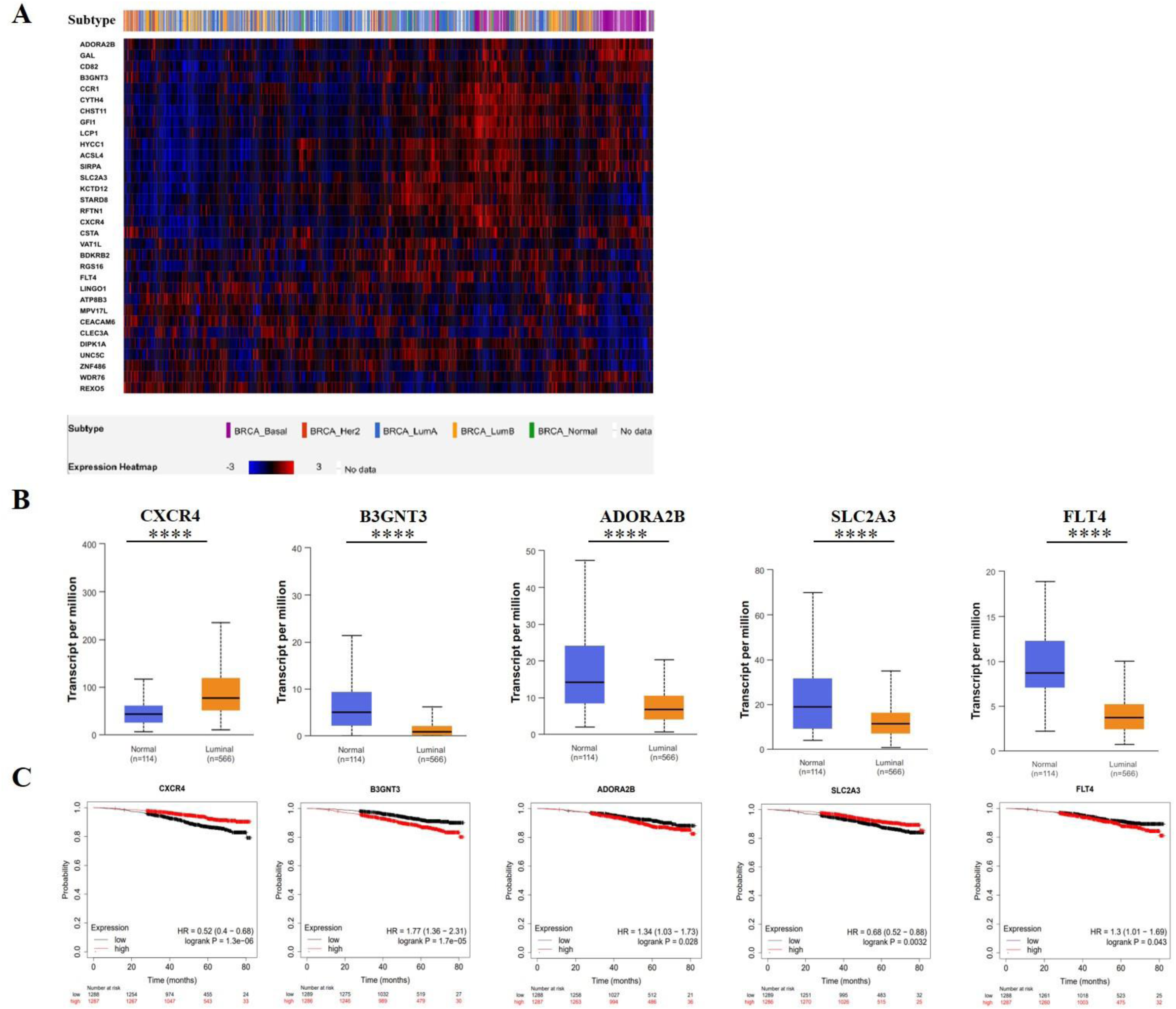
Gene expression in breast cancer patient samples. **(A)** Supervised hierarchical clustering analysis of 1084 samples (all subtypes) using mRNA expression z-scores relative to all samples (log RNA Seq. V2 RSEM) for a 73-gene ETR cassette. **(B)** Expression of upregulated genes of normal and primary tumor TCGA samples using the UALCAN database analysis (**** *p*<0.01). **(C)** Kaplan-Meier survival curves of select genes are shown.

We checked the expression of these 33 upregulated genes in the UALCAN database and the association of these genes with patient survival using KM plots (Figure 4 B, C and supplementary figure 2 B, C). As compared to normal, CXCR4, RGS16, and LCP1 were highly expressed in tumors but this indicated better overall patient survival. In contrast, B3GNT3, ADORA2B, SLC2A3, and FLT4 were less expressed in tumors when compared to normal samples but their high expression was associated with inferior overall survival. However, these genes were highly expressed in cells that showed ETR and in a sub-set of ER+ tumors from TCGA. It is tempting to suggest that increased re-expression of these 4 genes in tumor tissues during treatment-induced progression may indicate their escape from ET responsiveness. Testing the expression of these genes may be of value to determine an unfavorable prognosis during ET.

### 3.4. Identification of Differentially Expressed lncRNAs (DELs)

For the construction of CNC networks, we used the 7 up and 14 downregulated genes since they appeared in every significant pathway identified from DEGs and most were annotated as cell surface receptor proteins. If they were involved in ETR, they could be exploited as novel therapeutic targets. To construct CNC networks, we first estimated the expression of long non-coding RNAs in TAMR, LTEDI, and JOE cells (Figure 5 A, B). In JOE cells very few DELs were obtained (85 upregulated and 7 down-regulated) and very few of the lncRNAs overlapped with either TAMR or the LTEDI DELs. This suggests that JMJD6 may modulate the ETR cassette genes directly or by an alternate lncRNA mechanism distinct from TAMR-LTEDI. 959 DELs were identified in TAMR cells of which, 448 lncRNAs were upregulated and 511 lncRNAs were downregulated (adjusted p-value ≤ 0.05, log_2_FC=1) (Figure 5 A, B supplementary file 1). In LTEDI cells, 527 lncRNAs were differentially expressed, of which 198 were up and 329 were downregulated (adjusted p-value ≤ 0.05, log_2_FC=1) (Figure 5 A, B, supplementary file 1). Among DELs, 337 were common between TAMR and LTEDI cells and like the observation in DEGs, maintained the common directionality. 129 were upregulated and 200 were downregulated (Figure 5 B).

**Figure 5:**
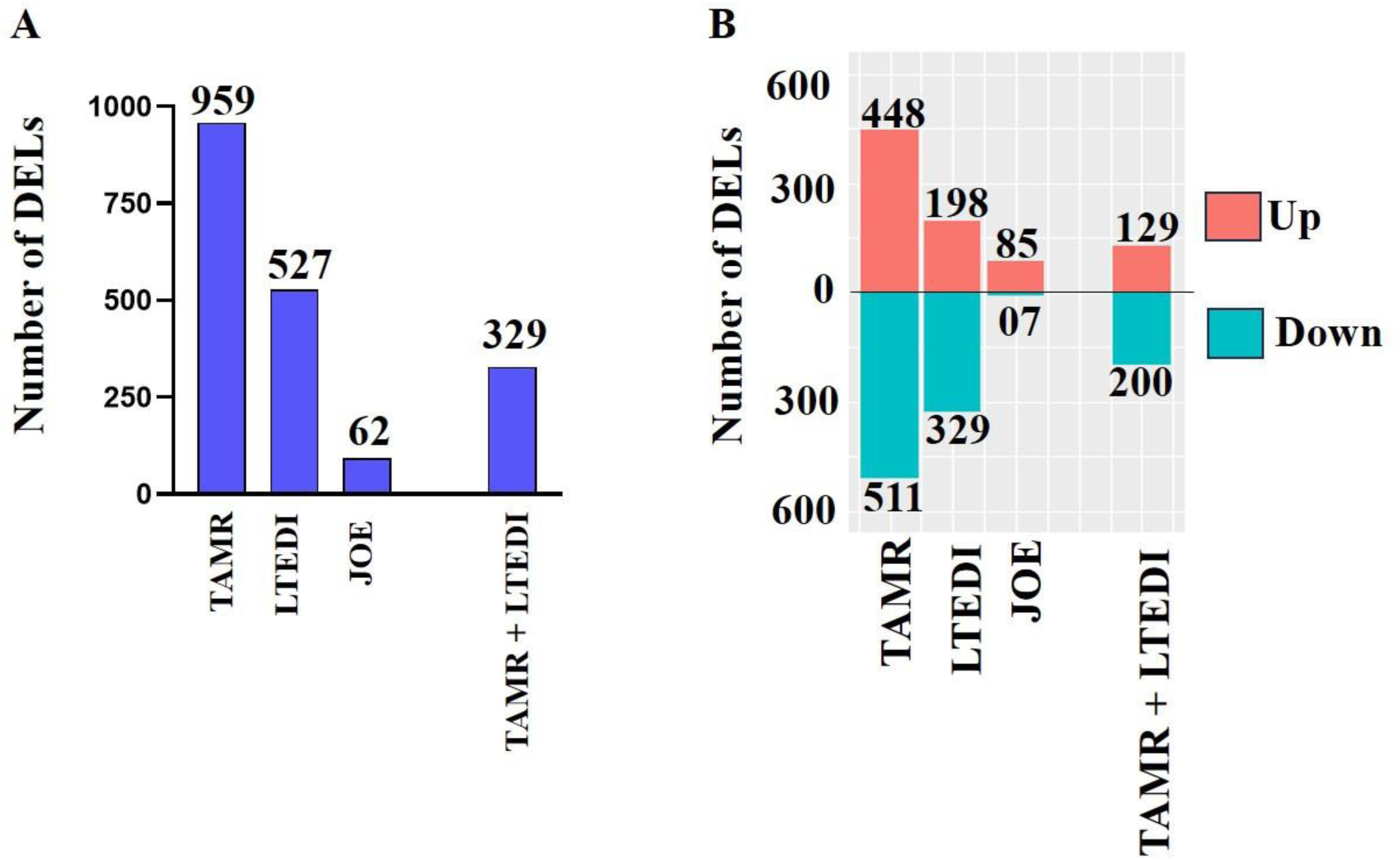
Estimation of DELs in three cell lines. **(A)** Differentially expressed long noncoding RNAs, **(B)** upregulated and downregulated long noncoding RNAs in TAMR, LTEDI, and JOE cell lines.

### 3.5. Construction of co-expression network

LncRNAs affect mRNA expression in both cis- and trans-regulatory manner. Positive correlation typically indicates cis-regulation of the mRNA by the lncRNA if the pairs are localized in close proximity to the genome (within 10 kb). To find such pairs, analysis of DEG:DEL pairs where both were upregulated, or both were downregulated was carried out by using Pearson correlation analysis, and positively correlated pairs were studied (Pearson correlation cut off > 0.98 for TAMR-LTEDI and > 0.95 for JOE). Four different DEG and DEL co-expression networks were constructed using the selected 7 up and 14 down DEGs in TAMR and LTEDI cells. 7 up DEGs positively correlated with 190 and 60 DELs in TAMR and LTEDI cells respectively (Supplementary figure 3 A, B). 14 downregulated DEGs had a positive correlation with 239 DELs in TAMR and 177 DELs in LTEDI (supplementary figure 3 C, D). As with DEGs, we were more interested in common rather than cell line-specific networks and identified a set of DELs that are commonly co-expressed in both TAMR and LTEDI. Specifically, 25 up-regulated DELs were co-expressed with 7 up DEGs (Figure 6 A), and 14 downregulated genes correlated with 93 common DELs (Figure 6 B). In JOE cells, only 5 DELs were common and only 3/7 upregulated genes had 55 positively correlated DELs, and 5/14 downregulated DEGs paired with 4 DELs (Figure 6 C, D). To establish cis-regulation, the genomic distance between the DEG:DEL pairs was calculated. None of the positively correlated mRNA-lncRNA pairs were placed near each other (within 10 kb), which suggested that the lncRNAs did not regulate the mRNAs via cis-regulation. Therefore, it is plausible that in ET resistance, cis-regulatory mechanisms play less of a role.

**Figure 6.**
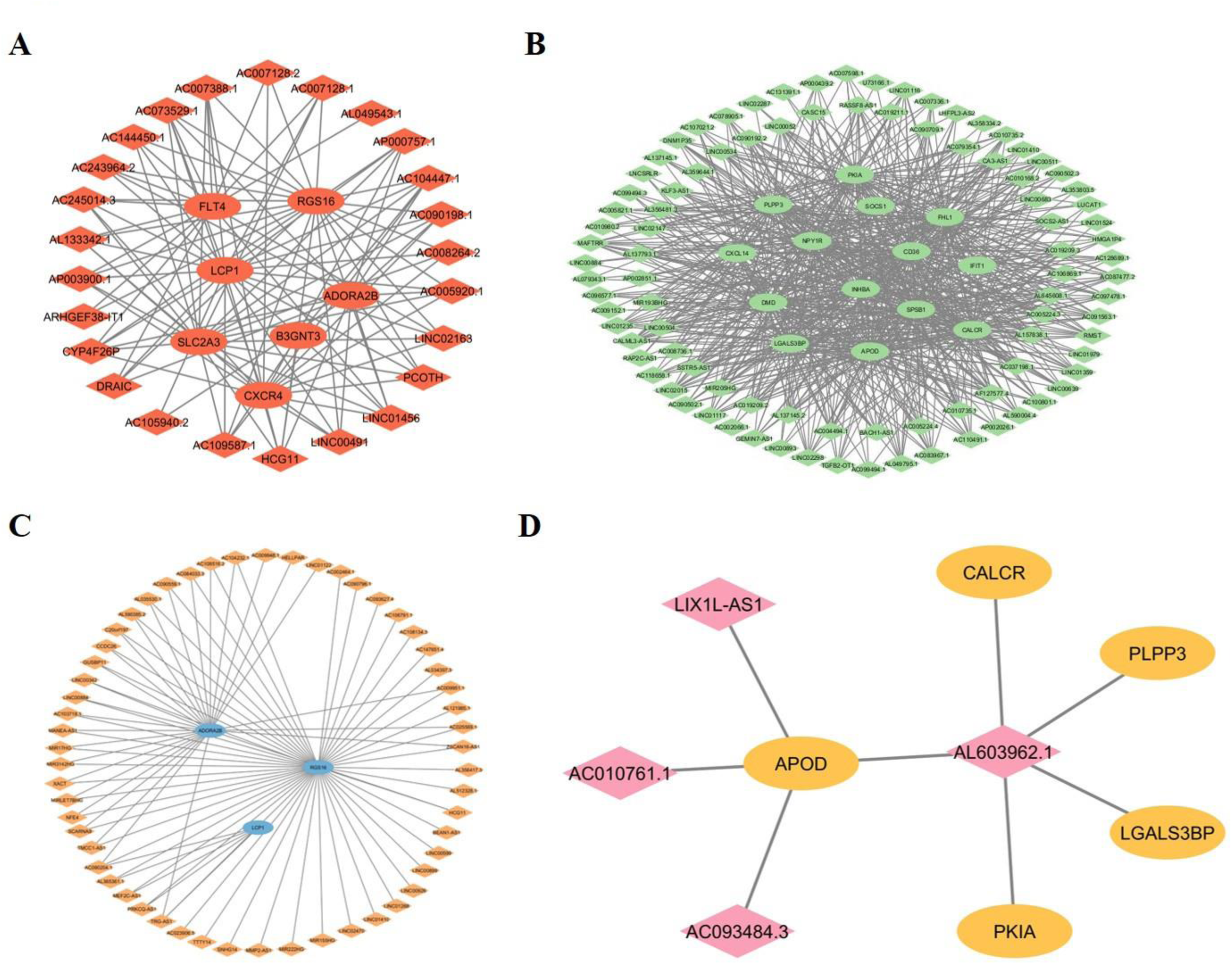
Construction of co-expression network. CNC network of TAMR and LTEDI cells using **(A)** 7 Upregulated DEGs correlated with 25 common DELs, **(B)** Downregulated DELs co-expressed with 7 upregulated DEGs. (B) 14 downregulated DEGs correlated with 93 common DELs. **(C)** Upregulated and **(D)** Downregulated DEL networks in JOE cells. The ellipse represents DEGs, and diamonds represent DELs.

### 3.6. Construction of competing endogenous RNA (ceRNA) network

Since lncRNAs act as an endogenous “sponge” to sequester microRNAs (miRNAs), thereby altering mRNA expression, we constructed ceRNA networks to explore the mRNA:lncRNA:miRNA connectivity. 21 DEGs (7 upregulated and 14 downregulated) from pathway-based curation were used. Since we did not have data for miRNA profiles for these cells, we used the following conditions to predict miRNA levels from the positively correlated DEG:DEL pairs-

1. if both lncRNA and mRNA from the CNC network are upregulated, it is likely that the miRNAs regulating them are depleted.
2. if both are low, miRNAs inhibiting them are likely upregulated.

Since TAMR-LTEDI cells showed a great degree of overlap but JOE had different DEG:DEL pairs, we made separate ceRNA networks. The TAMR/LTEDI/up network consisted of 15 miRNAs, 5 DEGs, and 5 DELs whereas, 11 miRNAs, 6 DEGs, and 8 DELs were found in the TAMR/LTEDI/down network (Figure 7 A and B respectively, Table 1). Together, 15 miRNAs are predicted to be downregulated and 11 miRNAs upregulated.

**Figure 7.**
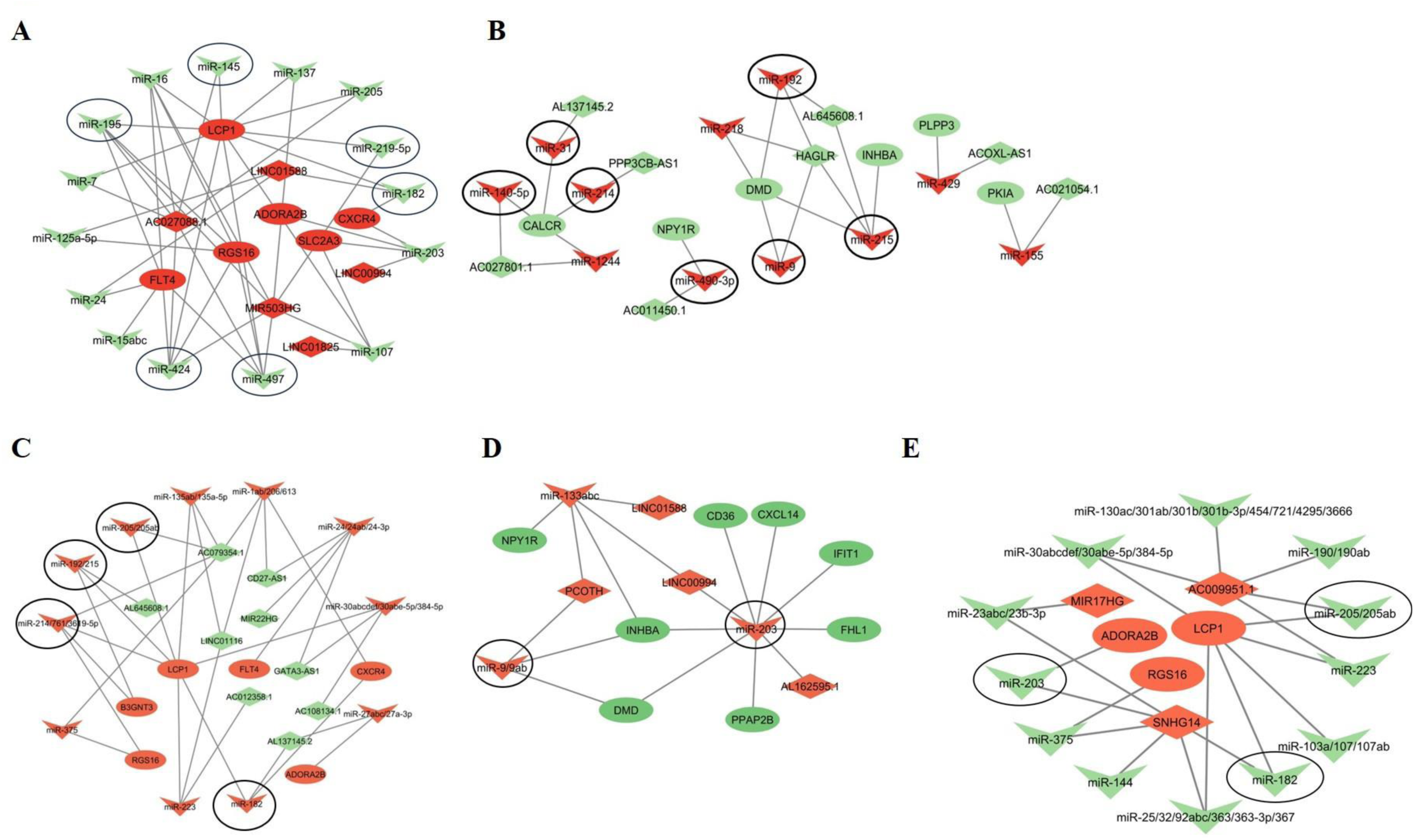
Construction of competing endogenous RNA (ceRNA) network using. **(A)** mRNAs (up), lncRNAs (up), and miRNAs (down) in the TAMR/LTEDI network, **(B)** mRNAs (down), lncRNAs (down), and miRNAs (up) in the TAMR/LTEDI network, **(C)** Network of upregulated mRNAs, miRNAs, and downregulated lncRNAs in TAMR and LTEDI cell lines, **(D)** mRNAs (down), lncRNAs (up), and miRNAs (up) in TAMR and LTEDI cell lines and **(E)** ceRNA network constructed in JOE cells using upregulated pairs. The ellipse represents mRNAs, the diamond represents lncRNAs, “V” shape represents miRNAs. For the prediction of miRNA levels-Red represents upregulated, and green/blue represents downregulated miRNAs.

**Table 1:**
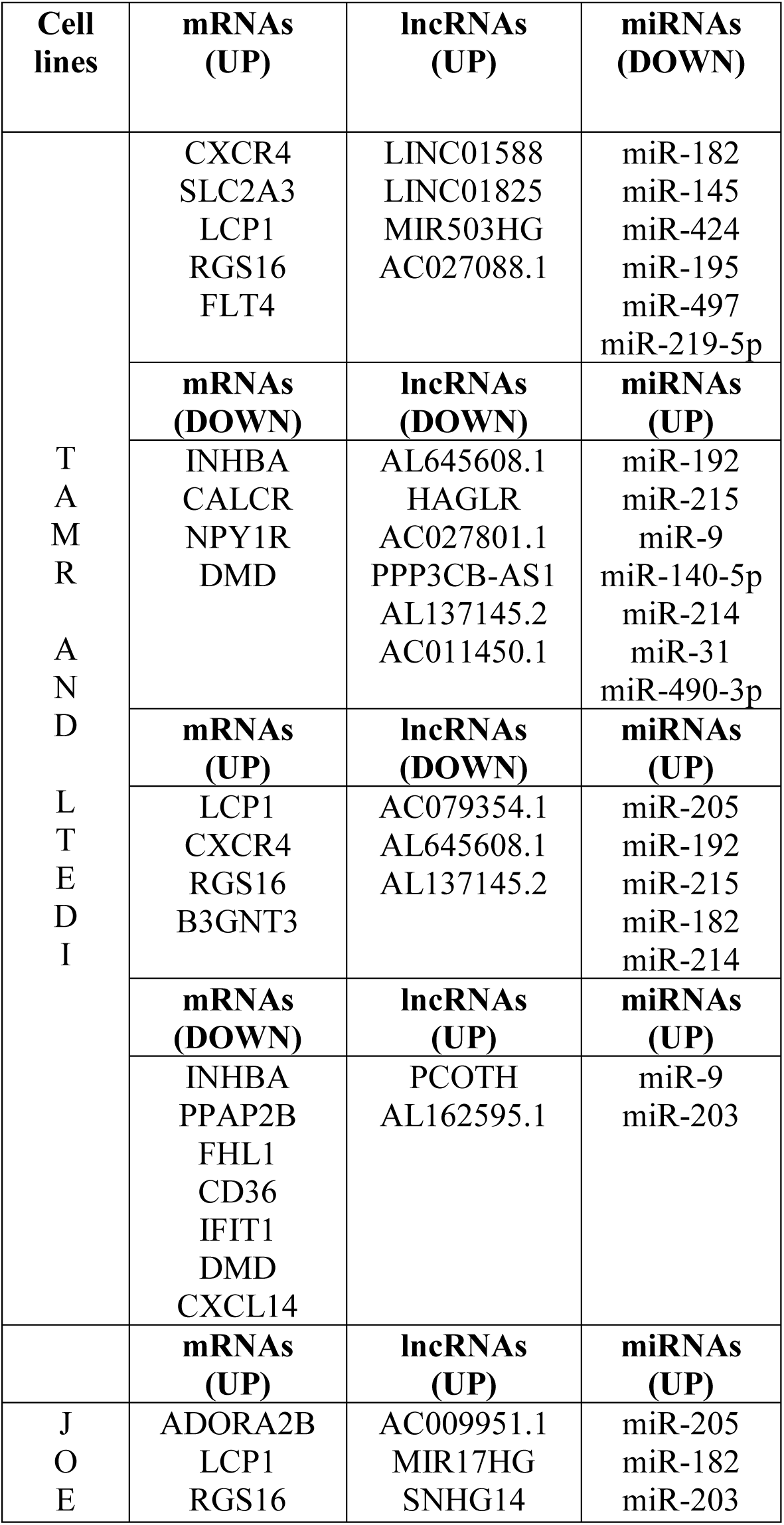
Triads in ETR cell line model systems.

Next, we identified negatively correlated DEG-DEL pairs (cut off > - 0.98). For negatively correlating DEG:DEL pairs,

1. miRNA binding could cause degradation of interacting lncRNAs or interacting mRNAs (26,27).
2. miRNAs binding to mRNAs may facilitate miRNA-driven upregulation of lncRNAs and vice-versa.
3. The negatively correlated network may represent a set of shared miRNAs that play a role in regulating both mRNAs and lncRNAs.

Figure 7 C shows the network of 6 upregulated DEGS in both TAMR and LTEDI paired with 9 downregulated DELs and 11 corresponding miRNAs. For 8 Downregulated DEGs common in both TAMR and LTEDI, 4 upregulated DELs and 3 miRNAs were obtained (Figure 7 D). From these studies, 14 miRNAs are predicted to be upregulated. In JOE cells, a separate ceRNA network was constructed using upregulated pairs, consisting of 12 miRNAs, 3 DEGs, and 3 DELs (Figure 7 E). We did not find any overlapping miRNAs for downregulated DEG-DEL pairs of JOE sets.

### 3.7. Prediction of miRNA status and validation

Since we did not have the miRNA expression data, we used ceRNA networks to predict the most likely expression level of the 40 interacting miRNAs in these cells. We predicted 25 to be upregulated and 15 to be downregulated. To validate our observations, we used publicly available miRNA expression data from GSE 56411 representing the regulation of miRNA in 3 different TAMR clones (28). Data from JOE and LTEDI cells was not available. 24/40 miRNAs were represented in the TAMR clone dataset and their ceRNA networks are shown in Supplementary figure 3. In 2 out of 3 TAMR clones, 7/11 miRNAs predicted to be upregulated and 6/13 predicted to be downregulated by our ceRNA network analysis showed the same directionality in expression (Supplementary Figure 5 A). Table 2 summarizes the mRNA-lncRNA-miRNA relationships for these miRNAs using our ceRNA networks. In particular, miRs-424/497/195 were found to be consistently downregulated in all 3 TAMR clones and these belong to the same family. We looked for common targets of these 3 miRNAs in our CNC networks and found FLT4 mRNA and the host lncRNA, MIR503HG. Expression of these miRNAs, FLT4 and MIR503HG were studied in ETR models by qRT-PCR (Figure 8). While the miRNAs were suppressed, both FLT4 and MIR503HG were highly expressed in ETR cells validating the RNA-seq data and miRNA expression prediction (Figure 8 A-E). Using UALCAN analysis of TCGA samples, miR-497, and miR-195 were found to be less expressed in tumors. In contrast, MIR503HG and miR-424 expression was higher in tumors when compared to normal samples (Supplementary Figure 5 B).

**Figure 8.**
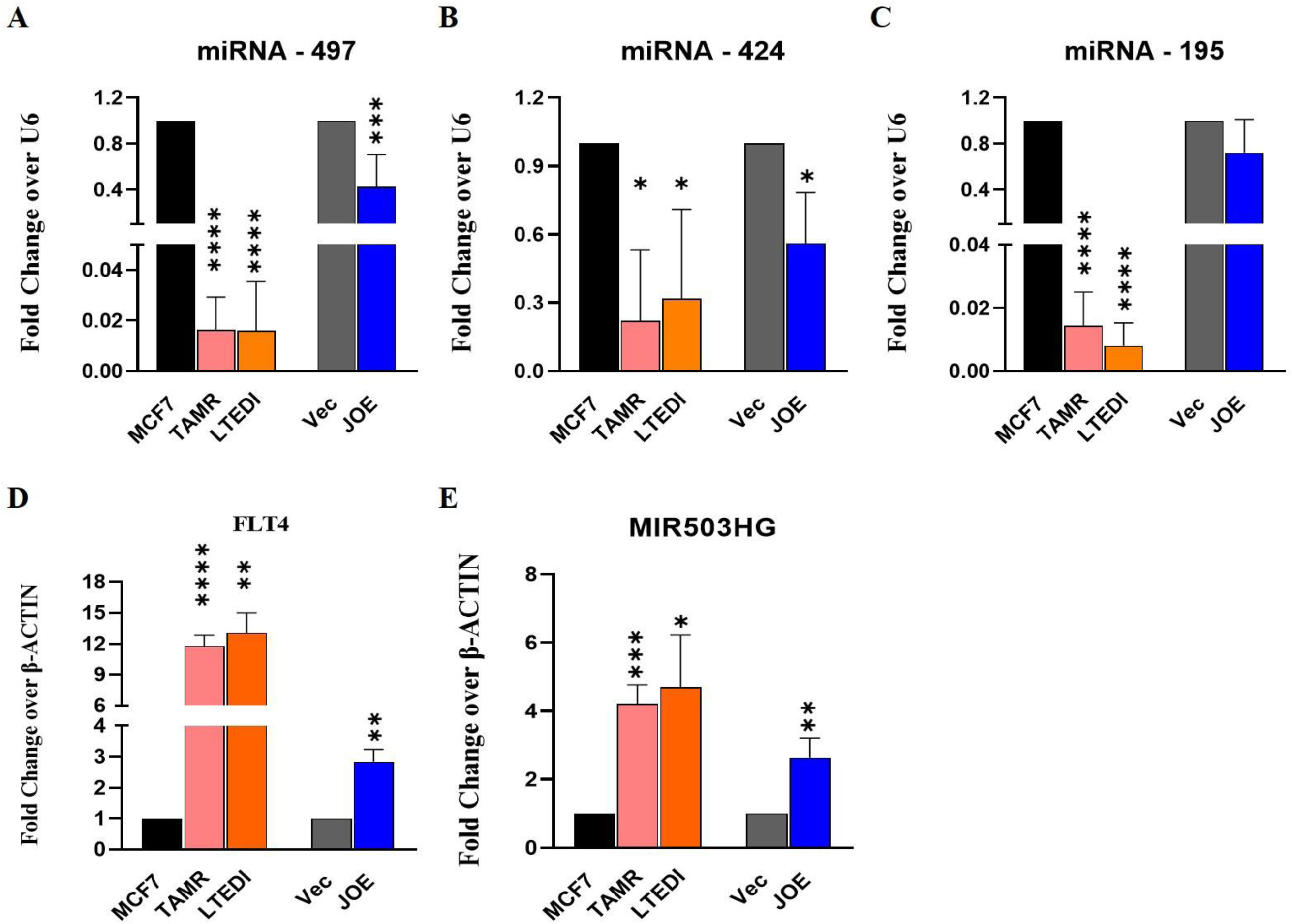
Validation of predicted miRNA expression. Real-time PCR analysis of miRNAs **(A-C)**, mRNA **(D),** and lncRNA **(E)** in ETR cell lines. Relative expression of miRNAs was normalized to a U6 and FLT4 and MIR503HG was normalized to beta-actin. MCF7 and “Vec” were used as respective controls and values were normalized to 1. *p ≤ 0.05, **p ≤ 0.01, and ***p ≤ 0.001.

**Figure 9:**
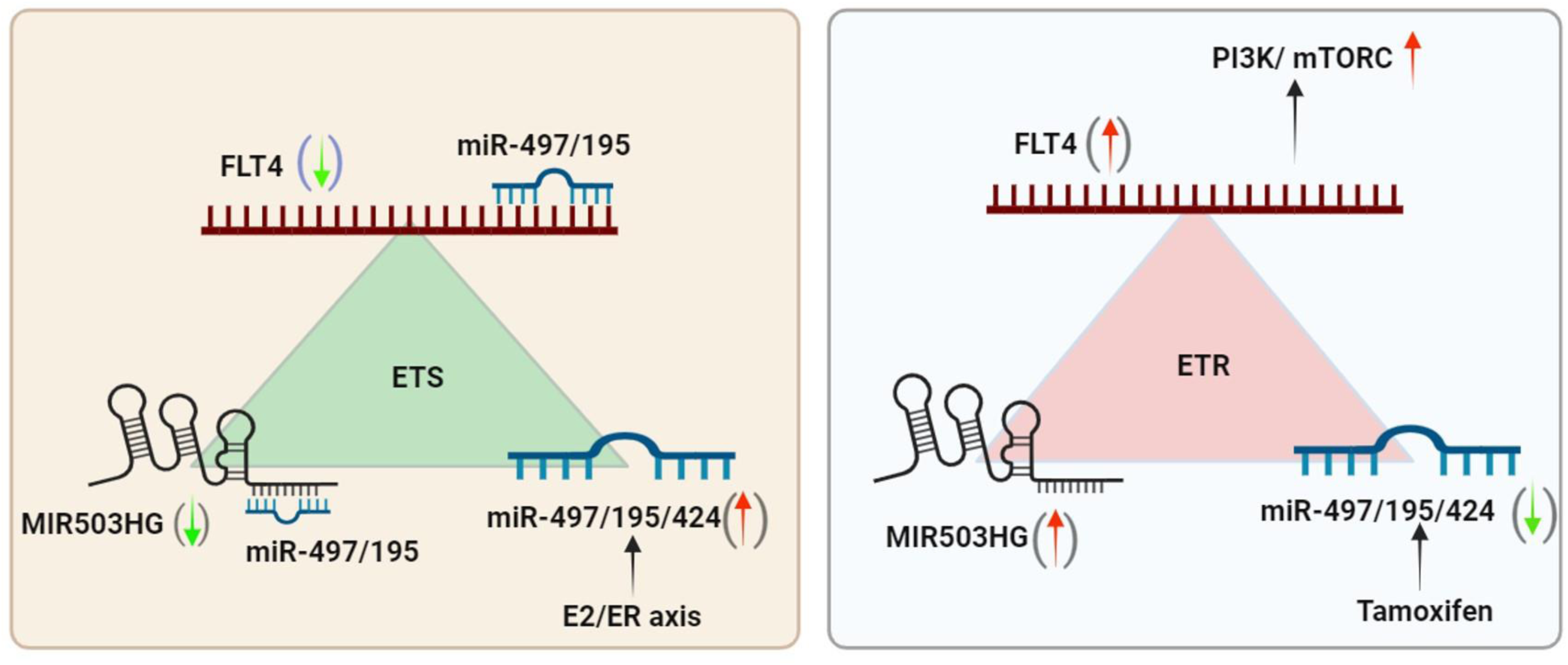
Gene expression in breast cancer patient samples. Graphical representation of the probable mechanisms of FLT4-MIR503HG-miR-497/195 in ETS (Endocrine Therapy sensitive) and ETR model systems.

**Table 2:**
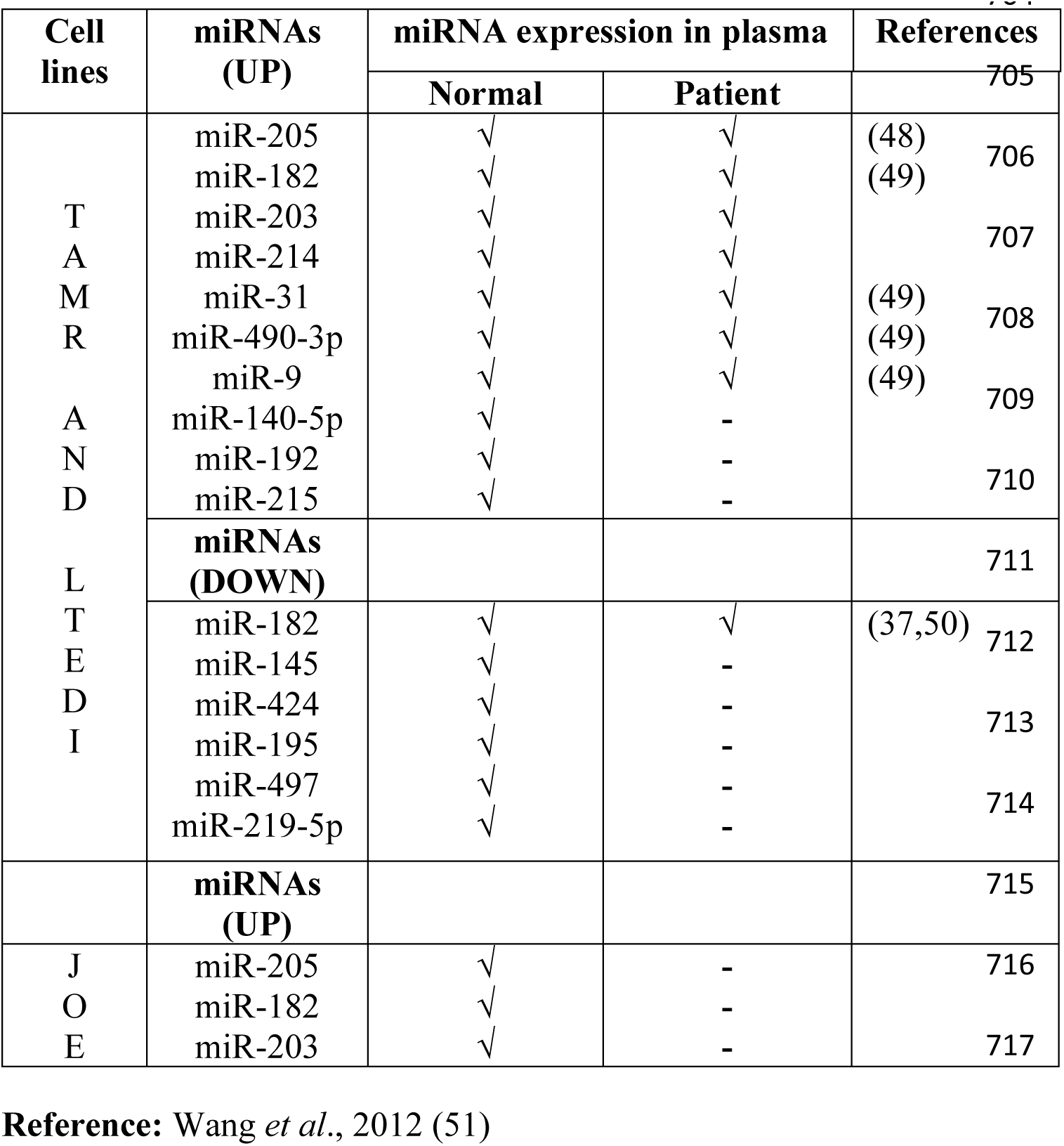
miRNA expression levels in the plasma sample.

## 4. DISCUSSION

Cells achieve ETR by different modes. Using TAMR/Fulvestrant resistant cells and ETR xenograft models, gene signatures have been developed but have not been deployed extensively in the clinical setting. Integration of data from other modes of ETR, such as lowering of ER expression or gain of agonistic activity in resistant cells is lacking in these gene panels. We hypothesized that combining information of RNA regulatory networks from cells representing distinct mechanisms of resistance may improve their clinical adaptability. We established and used three model systems TAMR, LTEDI, and JOE for transcriptomic analysis and subsequent CNC and ceRNA network construction. Of these, TAMR and LTEDI cells are the most used endocrine therapy model systems (29). Since in LTEDI cells, the Tam effect is controversial showing both agonistic and antagonistic activity, we first tested Tam effect on these cells. We report here that Tam behaves as a pure agonist allowing us to integrate 3 different modes of ETR in our studies.

RNA-Seq data of TAMR, LTEDI, and JOE cells identified 170 DEGs, of which, 73 (ETR cassette genes) were expressed in the same direction in all 3 cell lines. Pathway enrichment analysis identified crucial cancer hallmark pathways. For CNC and ceRNA analysis, we used pathway-based curation identifying 7 commonly upregulated and 14 commonly downregulated genes. Most genes were cell surface receptors and involved in various types of cancers (Table 3). CXCR4, B3GNT3, and ADORA2B are known markers of poor prognosis in breast cancer and therefore good targets for cancer treatment (30–32). These genes were tested in Fulvestrant-resistant cells that use increased EGFR and other tyrosine kinase member-driven ETR pathways, and 26 genes out of 85 were expressed in the same direction (33,34).

**Table 3:**
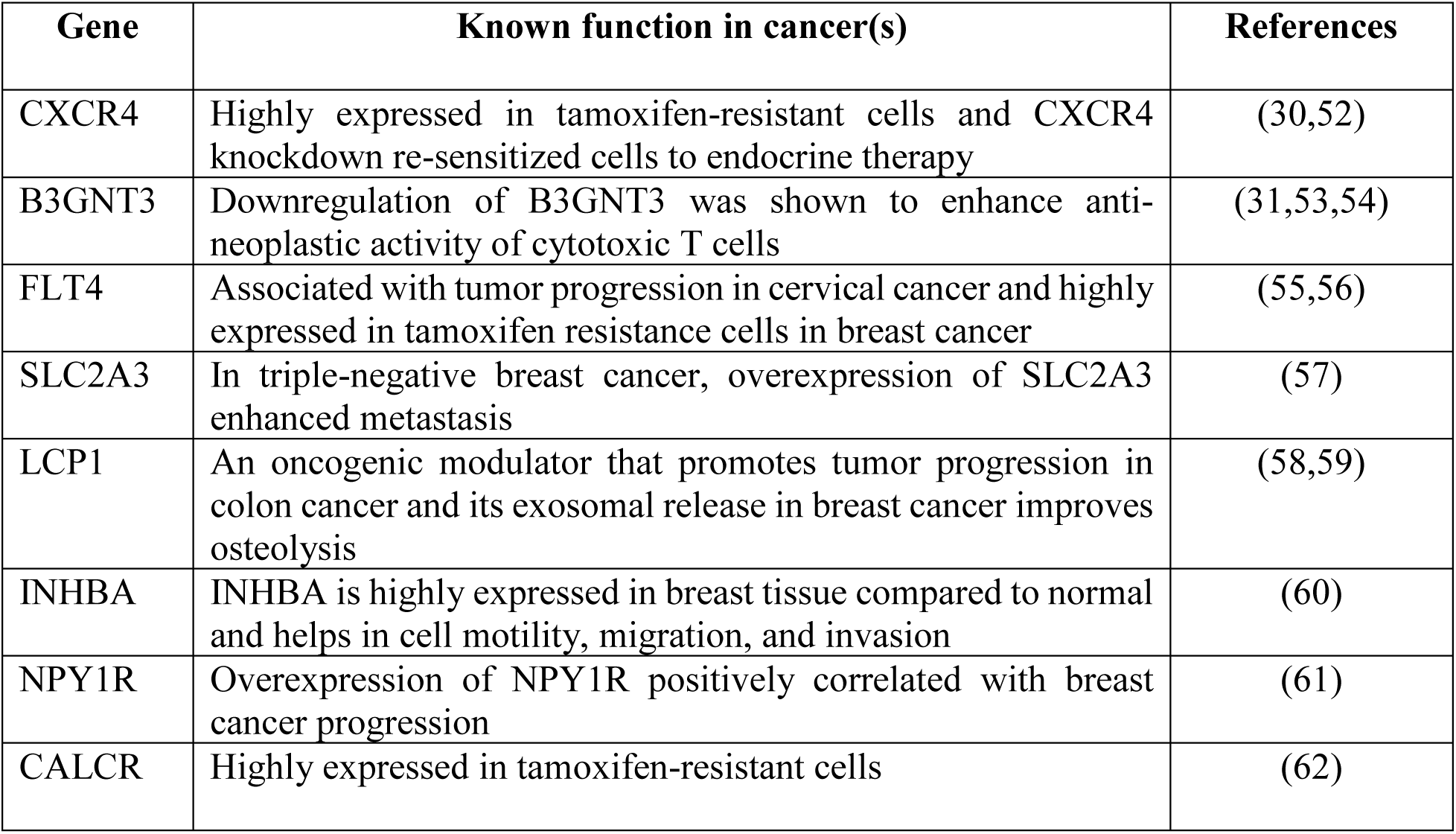
The seven upregulated genes and their role in cancers.

| Gene | Known function in cancer(s) | References |
| --- | --- | --- |
| CXCR4 | Highly expressed in tamoxifen-resistant cells and CXCR4 knockdown re-sensitized cells to endocrine therapy | (30,52) |
| B3GNT3 | Downregulation of B3GNT3 was shown to enhance anti-neoplastic activity of cytotoxic T cells | (31,53,54) |
| FLT4 | Associated with tumor progression in cervical cancer and highly expressed in tamoxifen resistance cells in breast cancer | (55,56) |
| SLC2A3 | In triple-negative breast cancer, overexpression of SLC2A3 enhanced metastasis | (57) |
| LCP1 | An oncogenic modulator that promotes tumor progression in colon cancer and its exosomal release in breast cancer improves osteolysis | (58,59) |
| INHBA | INHBA is highly expressed in breast tissue compared to normal and helps in cell motility, migration, and invasion | (60) |
| NPY1R | Overexpression of NPY1R positively correlated with breast cancer progression | (61) |
| CALCR | Highly expressed in tamoxifen-resistant cells | (62) |

We constructed a CNC network using DEGs and DELs in both TAMR-LTEDI cells and separately in JOE cells. We failed to find cis-regulation of mRNAs by lncRNAs as a mechanism for co-expression and suggest that this may not be a preferred regulatory mechanism in ETR. The most relevant ceRNA networks were obtained from the pathway-based curation, that is 7 induced and 14 suppressed genes. 27 lncRNAs were found using both positively and negatively correlated DEG:DEL pairs. Since, we did not have miRNA expression profiles, we predicted miRNA expression based on the ceRNA hypothesis. We obtained a total of 40 miRNAs in this network and found 24 of them in published TAMR miRNA data (28). Of these, we could correctly predict the expression level of 7/11 upregulated and 6/13 downregulated miRNAs. A family of miRNAs-miR-424/195/497 was consistently lower in these TAMR clones. Using our CNC and ceRNA networks we identified FLT4 and MIR503HG as their targets and RNA-seq data indicated that they were highly expressed in ETR models. Vilquin *et al* have shown that depletion of miR-424 and upregulation of miR-205 resulted in Aromatase Inhibitor resistance phenotype in ER+ cells (35). miR-497-5p and miR-195 interfere with PI3K-mTOR signalling and MIR503HG is known to sponge the interfering miRNAs to promote subsequent ETR (36). These miRNAs are embedded in lncRNA MIR497HG and their downregulation was associated with TAM resistance (37). Both miRNAs contain ER binding sites in their promoter and they are upregulated by E_2_ treatment so in the presence of Tam, their expression would be abrogated. MIR497HG expression was also high in ETR cells. We have used an exceptionally high cut off (r^2^ =0.98) for our CNC networks and hence it was not represented in our final list. In addition, miR-424 arises from MIR503HG and its reduced expression is known to increase the expression of VEGF for tumor neo-angiogenesis (36,38). Based on these observations, we chose to validate the expression of the FLT4:MIR503HG:miR-424/497/195 triad by quantitative RT-PCR. The miRNAs were downregulated and we suggest that this caused the higher expression of their targets, FLT4 and MIR503HG. In JOE cells, we obtained very few DELs. Nevertheless, we performed quantitative RT-PCR in these cells and show patterns of expression similar to TAMR-LTED cells. Whether these patterns are perpetuated by MIR503HG based sponging of identified miRNAs or due to the loss of a functional E2/ER axis, requires additional experimentation. Further, its possible that JMJD6 uses the depletion of ER as the major ETR mechanism rather than altering ceRNA networks. In contrast, presence of abundant ER expression in TAMR and LTEDI may provoke a more complex RNA regulation of ETR mechanisms in these cells.

FLT4 is kinase and a receptor for VEGF and is usually expressed in endothelial cells leading to increased lymph-angiogenesis based metastasis. However, its increased expression in epithelial cells is shown to increase the proliferation, motility, and invasion of breast cancer cells (39). Higher expression of the FLT4-VEGF-C axis promotes invasion and metastasis in cancer and interfering with it decreases cell proliferation (40,41). Together, the network of FLT4, MIR503HG, and miR-424/497/195 indicate the abolishment of E_2_ activity, augmentation of PI3K-mTOR signalling, and aggressive behaviour of cancer cells. In ETR, high FLT4 therefore makes for a good therapeutic target (42).

Most miRNAs identified in this study can be quantified in both normal healthy individuals as well as in miRNA-based signatures in cancer patients via liquid biopsy approaches (Table 2) (43). Curiously, none of the ETR cassette genes have been described previously in TAMR signature genes (8,44,45), nor did they overlap with the EndoPredict (3 proliferation and 5 hormone receptor-related genes from Myriad Genetics), nor with the ENDORSE prognostic model for ET that identified MYC targets, E2F1 targets, and PI3KC-mTOR signaling pathway gene expression in tumors as indicators of high risk for ETR (46,47). Our network suggests that these pathways from ENDORSE could be represented in an alternate manner via the RNA networks encompassing small and long non-coding RNAs. Together, these data significantly validate the usefulness of the ceRNA network-based approach to predict actionable miRNAs.

We next checked the status of the ETR cassette genes in TCGA datasets. We hypothesized that if ER+ tumors were to be resistant to ET, they should display certain gene expression profiles like ER-tumors as they no longer use the E2/ER axis for sustenance. When we assessed the expression of ETR cassette genes in TCGA breast cancer data, the expression pattern of upregulated genes led to the clustering of a subset of ER+ tumors with ER-negative breast cancers (Figure 4). This suggests that genes appearing in ETR cell data can segregate the ER+ tumors. Next, to determine the individual profile of each gene, we employed the UALCAN database. CXCR4, RGS16, and LCP1 displayed higher expression in tumors when compared to normal breast but did not associate with poor outcomes in ER+ breast cancer. Interestingly, B3GNT3, ADORA2B, SLC2A3, and FLT4 expression was lower in tumors and higher in normal breast, and low expression was associated with better survival outcomes. This suggests that the re-expression of these genes may indicate the onset of ETR. In tumors, TCGA data showed that the miR-195/497 are suppressed in breast cancer samples biopsied at diagnosis and possibly re-expression of FLT4 may not be governed by miRNA expression alone. However, one possibility is that FLT4 expression switches from endothelial cells to tumor epithelium during the establishment of ETR. Future immunohistochemical studies are required to confirm this hypothesis.

## 5. CONCLUSION

Our discovery of FLT4:MIR503HG:miR-424/497/195 triad showed that in the absence of miRNA profiling data, total RNA-seq and RNA network analysis can predict actionable miRNAs. Based on our RNA regulatory networks in ETR cells, we suggest that in endocrine therapy sensitive (ETS) cells, miR-424/497/195 expression is high leading to suppression of FLT4 and MIR503HG. Re-expression of FLT4 mRNA, high expression of MIR503HG in tumors, probably occurs due to suppression of miR-497, miR-195, and miR-424 leading to the onset of ETR in cancer (Figure 8).

## 6. ABBREVIATIONS

AIs: Aromatase Inhibitors, ET: Endocrine Therapy, SERDs: Selective Estrogen Receptor Degraders, SERMs: Selective Estrogen Receptor Modulators, Tam: Tamoxifen, MBC: Metastatic Breast Cancer, JMJD6: Jumonji Domain Containing 6, JOE: JMJD6 overexpressing MCF7 cells, TAMR: Tamoxifen resistant cells, LTEDI: Long-Term Estrogen Deprived-Independent cells, RNA-seq: RNA sequencing, ceRNAs: competing endogenous RNAs, lncRNA: Long non-coding RNA, miRNA: microRNA, 4-OHT: 4-hydroxy Tamoxifen, DEGs: Differentially Expressed Coding Genes, DELs: Differentially Expressed long non-coding RNAs.

## 9. AUTHOR CONTRIBUTIONS

**Saheli Pramanik:** Methodology, Data curation, Visualization, **Partha Das**: Investigation, Visualization, Experimental data **Monalisa Mukherjee**: Visualization, Investigation. **Kartiki V Desai** Conceptualization, Analysis, Supervision. **All authors** contributed to Writing-Original draft preparation, reviewing, and editing the manuscript.

## ACKNOWLEDGEMENT

The authors would like to thank Aritra Gupta and Siddarth Bharadwaj for their technical help and assistance in editing the manuscript.

---

**Supplementary figure 1:**
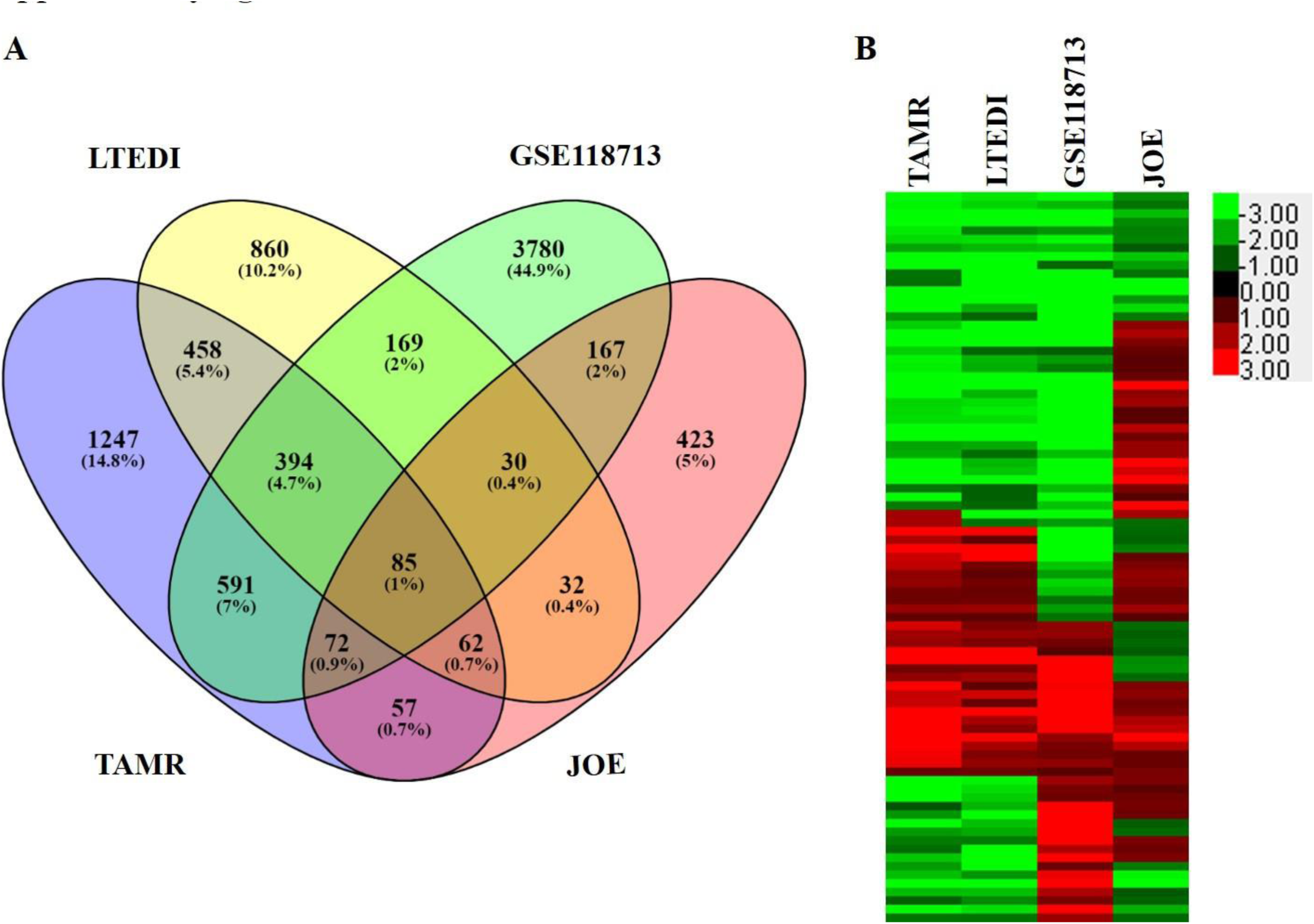
RNA-seq analysis of Fulvestrant data (GSE118713). **(A)** Venn diagram of the common genes between TAMR, LTEDI, JOE, and GSE118713. **(B)** Expression patterns of 85 genes were shown by heatmap. Changes in expression levels were displayed from green (downregulated) to red (upregulated), as shown in the color gradient on the right side.

**Supplementary figure 2:**
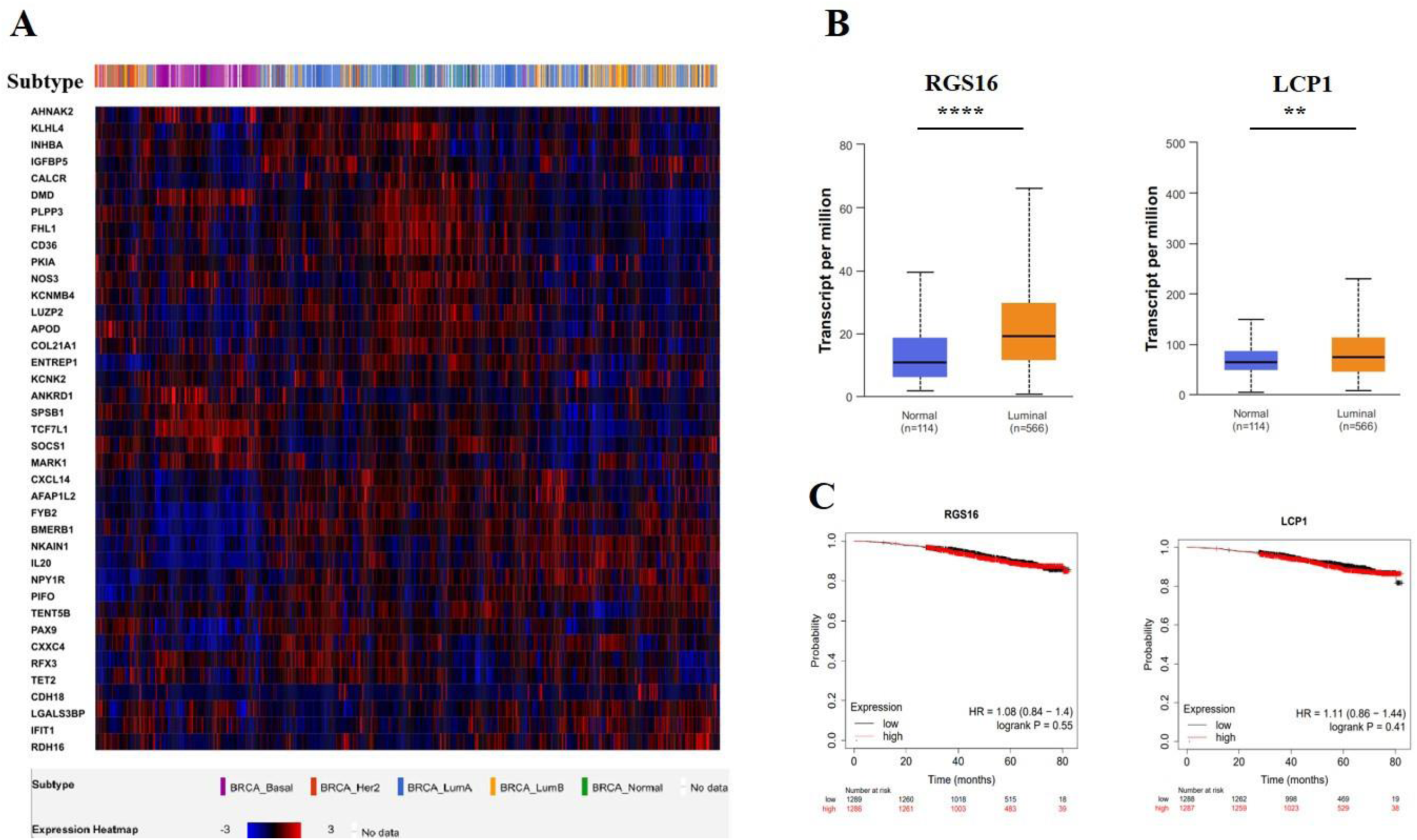
**Gene expression in breast cancer patient samples from the publicly available dataset**. **(A)** Supervised hierarchical clustering analysis of 1084 samples (all subtypes) using mRNA expression of downregulated genes (log RNA Seq. V2 RSEM). **(B)** Expression of upregulated genes of normal and primary tumor TCGA samples using the UALCAN database analysis (**** P<0.01). **(C)** Kaplan-Meier survival plot of select genes.

**Supplementary figure 3:**
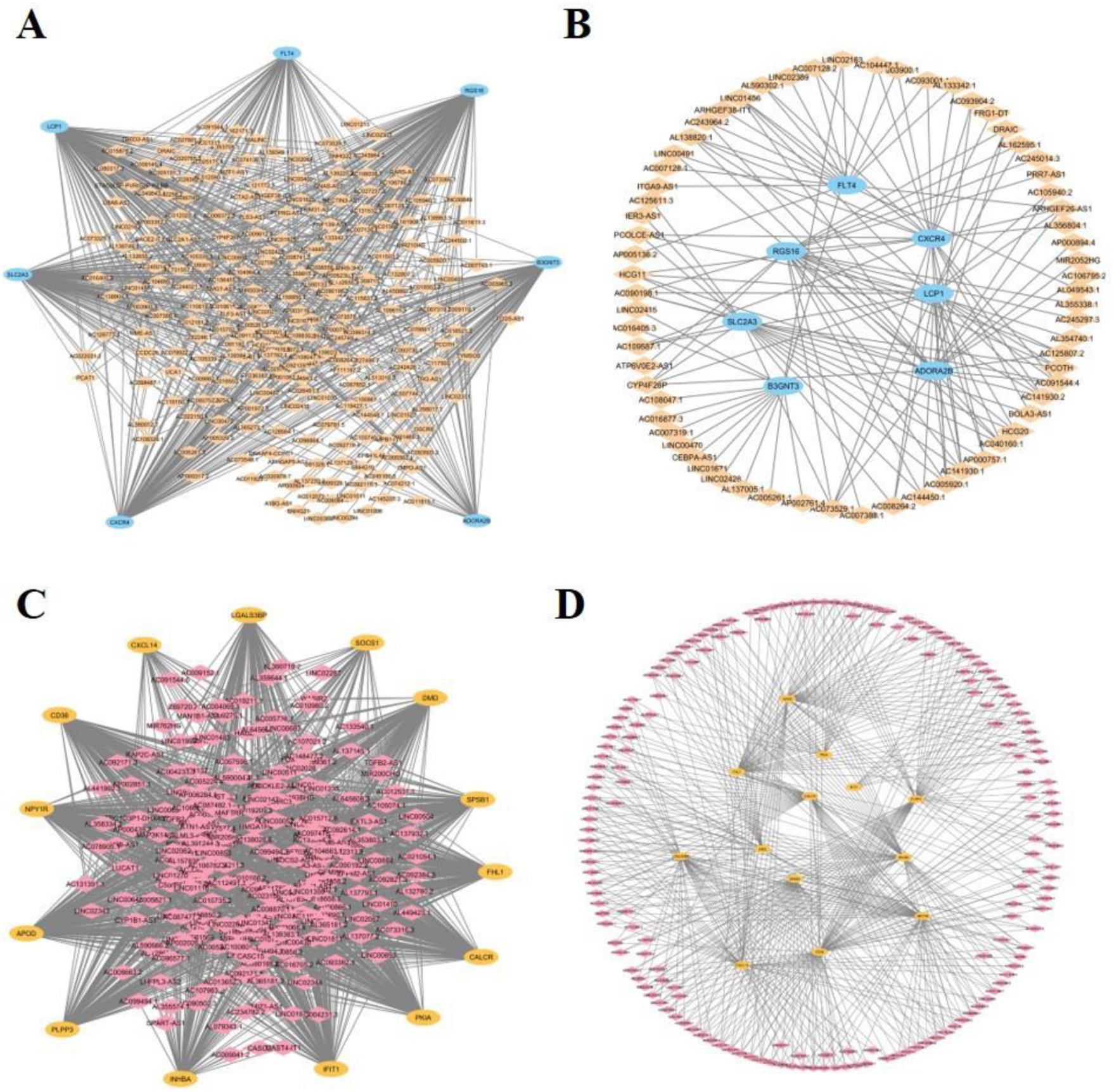
CNC network of individual TAMR and LTEDI cells. **(A and B)** upregulated DEGs:DELs and **(C and D)** downregulated DEGs:DELs in TAMR and LTEDI cells respectively.

**Supplementary figure 4:**
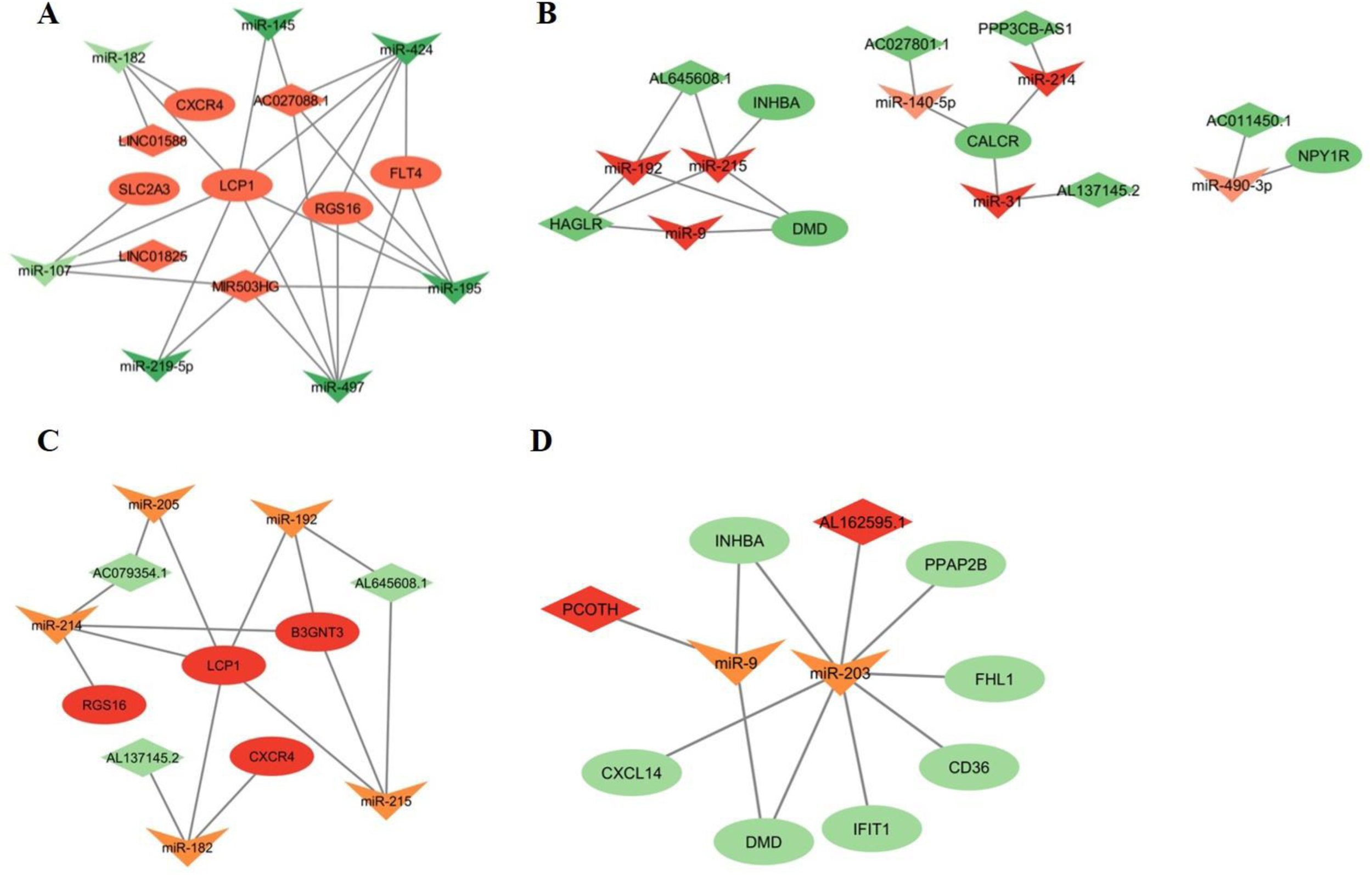
Construction of competing endogenous RNA (ceRNA) network. The upregulated ceRNA network of TAMR-LTEDI cells. **(A)** upregulated, **(B)** downregulated. **(C)** ceRNA network of upregulated DEGs and downregulated DELs from TAMR and LTEDI cells. **(D)** ceRNA network of downregulated DEGs and upregulated DELs from TAMR and LTEDI cells. The ellipse represents mRNAs, the diamond represents lncRNAs, “V” shape represents miRNAs.

**Supplementary figure 5:**
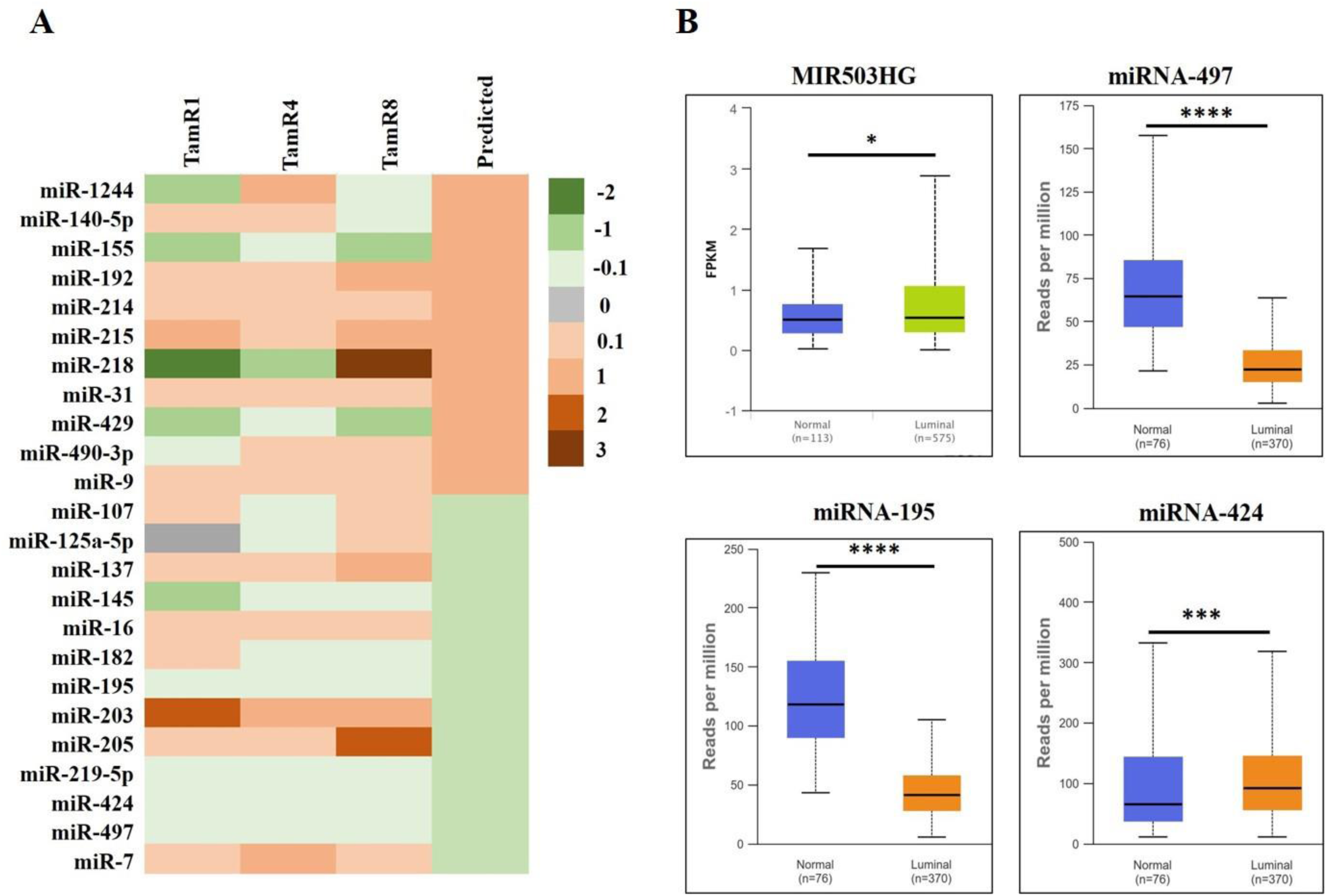
**(A) Publicly available miRNA expression data from GSE56411 from 3 different TAMR clones compared with the prediction of miRNA regulation from this study**. Validated miRNAs were marked by a circle in Figure 7. **(B)** Expression of MIR503HG, and miR-497/195/424 in normal and tumor TCGA samples derived from UALCAN database analysis (**** p<0.01).

**Supplementary file 1:** List of DEGs and DELs identified in TAMR and LTEDI cells (adjusted p≤0.05)

**Supplementary table 1:** Primers used for qRT-PCR analysis.

