## Supplementary Table 1 for "Suppression of miRs-497/195 axis possibly confers endocrine therapy resistance via elevated expression of FLT4 and the noncoding RNA MIR503HG"

| **Gene_Name** | **Primer Sequence 5' - 3'** |
| --- | --- |
| **FLT4-RT-F** | TGCGAATACCTGTCCTACGATGC |
| **FLT4-RT-R** | TCCTGGTGCAAGTTTTGAAAAT |
| **MIR503HG-F** | TCCCGCCAAATGAGTCAGTC |
| **MIR503HG-R\** | CCTGTGTGGGGTTCCACTTT |
| **Beta-actin F** | GAG CAC AGA GCC TCG CCT TT |
| **Beta-actin R** | TCA TCA TCC ATG GTG AGC TGG |
| **U6-RT-F** | CTCGCTTCGGCAGCACATATACT |
| **U6-RT-R** | ACGCTTCACGAATTTGCGTGTC |
| **miR-424 ST-LOOP** | GTCGTATCCAGTGCAGGGTCCGAGGTATTCGCACTGGATACGACAAACTT |
| **miR-497 ST-LOOP** | GTCGTATCCAGTGCAGGGTCCGAGGTATTCGCACTGGATACGACCAAACA |
| **miR-195 ST-LOOP** | GTCGTATCCAGTGCAGGGTCCGAGGTATTCGCACTGGATACGACTAACCG |
| **miR-424-F** | CACGCACAGCAGCAATTC |
| **miR-497-F** | CACGCACAGCAGCACACT |
| **miR-195-F** | CACGCATAGCAGCACAGA |
| **miRNA reverse primer** | GTGCAGGGTCCGAGGT |
